# Genomic characterisation and conservation genetics of the indigenous Irish Kerry cattle breed

**DOI:** 10.1101/210229

**Authors:** Sam Browett, Gillian McHugo, Ian W. Richardson, David A. Magee, Stephen D. E. Park, Alan G. Fahey, John F. Kearney, Carolina N. Correia, Imtiaz A. S. Randhawa, David E. MacHugh

## Abstract

Kerry cattle are an endangered landrace heritage breed of cultural importance to Ireland. In the present study we have used genome-wide SNP data (Illumina^®^ BovineSNP50 array) to evaluate genomic diversity within the Kerry cattle population and between Kerry cattle and other European cattle breeds. Visualisation of patterns of genetic differentiation and gene flow among cattle breeds using phylogenetic trees with ancestry graphs highlighted, in particular, historical gene flow from the British Shorthorn breed into the ancestral population of modern Kerry cattle. Principal component analysis (PCA) and genetic clustering emphasised the genetic distinctiveness of Kerry cattle relative to comparator British and European cattle breeds. Modelling of genetic effective population size (*N*_e_) revealed a demographic trend of diminishing *N*_e_ over time and that recent estimated *N*_e_ values for the Kerry breed may be less than the threshold for sustainable genetic conservation. In addition, analysis of genome-wide autozygosity (*F*_ROH_) showed that genomic inbreeding has increased significantly during the 20 years between 1992 and 2012. Finally, signatures of selection revealed genomic regions subject to natural and artificial selection as Kerry cattle adapted to the climate, physical geography and agro-ecology of southwest Ireland.

**Note 1:** This is an Associate Editor (D.E.M) Inaugural Article submission to *Frontiers in Genetics: Livestock Genomics*

**Note 2:** British English language style preferred for publication of this article.

## 1. Introduction

Approximately 10,000 years ago, humans first domesticated wild aurochs (*Bos primigenius*)—the progenitor of modern cattle—in the Fertile Crescent region of Southwest Asia (Edwards et al., 2010; Larson and Fuller, 2014; Larson et al., 2014; Park et al., 2015; MacHugh et al., 2017). Extant domestic cattle, which encompass humpless taurine (*B. taurus*), humped zebu (*B. indicus*) and myriad *B. taurus/indicus* hybrid populations, have, through genetic drift and natural and artificial selection, diversified into more than 1,100 recognised breeds. However, starting in the middle of the 20^th^ century, socioeconomic preferences for large highly productive dairy, beef and dual-purpose breeds have led to extinction and increased vulnerability of more than 200 locally-adapted landrace or native cattle breeds (Gandini et al., 2004; Food and Agriculture Organization, 2007; 2015).

With the advent of accelerating climate change, particularly in the Arctic and circumarctic regions (Vihma, 2014; Gao et al., 2015), agro-ecological environments in north-western Europe will inevitably undergo significant change during the coming century (Smith and Gregory, 2013; Wheeler and von Braun, 2013). It is, therefore, increasingly recognised that long-term sustainability of animal production systems and food security will necessitate conservation and management of livestock genetic resources in this region (Hoffmann, 2010; Boettcher et al., 2015; Kantanen et al., 2015). Locally-adapted native livestock breeds with distinct microevolutionary histories and minimal external gene flow will have accumulated novel genomic variation and haplotype combinations for quantitative health, fertility and production traits (Hill, 2014; Felius et al., 2015; Kristensen et al., 2015). These populations may therefore be key to future breeding programmes directed towards adaptation of European livestock to new agro-ecological and production environments (Biscarini et al., 2015; Boettcher et al., 2015; Phocas et al., 2016a; b).

The availability of powerful and cheap tools for genotyping large numbers of single nucleotide polymorphisms (SNPs) has provided conservation biologists and animal geneticists with the opportunity to characterise genomic variation and estimate population genetic parameters at very high resolution in threatened or endangered livestock breeds (Pertoldi et al., 2014; Ben Jemaa et al., 2015; Beynon et al., 2015; Meszaros et al., 2015; Burren et al., 2016; Decker et al., 2016; Iso-Touru et al., 2016; Manunza et al., 2016; Mastrangelo et al., 2016; Visser et al., 2016; Williams et al., 2016; Francois et al., 2017). These studies are already providing important baseline data for genetic conservation and will underpin programmes for managed breeding and biobanking of these populations (Groeneveld et al., 2016).

As a native breed with a claimed ancient heritage, Kerry cattle are considered culturally important to Ireland (Curran, 1990). It is a landrace cattle population that remains productive in harsh upland regions with poor quality feed, which are typical of southwest Ireland where the Kerry breed evolved (Food and Agriculture Organization, 2017). These cattle were often referred to anecdotally in Ireland as the “poor man’s cow” due to their ability to produce relatively large quantities of milk on very sparse fodder; the Kerry breed is also considered to be a remnant of what was once a substantially larger and more widespread historical population. Levels of inbreeding have been estimated using pedigree data and the accumulated figure since the foundation of the herd book in 1887 reached 15% in 1985 (O'hUigín and Cunningham, 1990).

In recent decades the Kerry cattle breed has experienced significant population fluctuations due to changing socioeconomic and agricultural circumstances. During the 1980s, the number of breeding females decreased to less than 200, prompting the Irish agricultural authorities to introduce a Kerry cattle conservation scheme (McParland, 2013), which has continued to the present day in the form of the Department of Agriculture, Food and the Marine (DAFM) Kerry Cattle Premium Scheme (Department of Agriculture Food and the Marine, 2017).

The formal conservation policy and supports initiated during the early 1990s led to a significant increase in the Kerry cattle population, such that by 2007 the number of breeding females had increased to more than a thousand animals (Food and Agriculture Organization, 2017). In recent years, however, due to deteriorating economic circumstances in Ireland post-2008, the Kerry cattle population has substantially declined once again and is classified as endangered and under significant threat of extinction or absorption through crossbreeding with other breeds (McParland, 2013; Department of Agriculture Food and the Marine, 2014).

The Kerry cattle breed was one of the first European heritage cattle breeds to be surveyed using molecular population genetics techniques. We have previously used autosomal microsatellite genetic markers and mitochondrial DNA (mtDNA) control region sequence variation for comparative evolutionary studies of genetic diversity in Kerry cattle and other British, European, African and Asian breeds (MacHugh et al., 1997; MacHugh et al., 1998; MacHugh et al., 1999; Troy et al., 2001). In addition, Bray and colleagues have used microsatellites to examine admixture and ancestry in Kerry cattle and the Dexter and Devon breeds (Bray et al., 2009). Results from these studies demonstrated that Kerry cattle exhibit markedly low mtDNA sequence diversity, but autosomal microsatellite diversity comparable to other cattle breeds native to Britain and Ireland. More recently, analyses of medium-and high-density SNP genotypes generated using genome sequence data from an extinct British *B. primigenius* subfossil have shown that Kerry cattle retain a significant genomic signature of admixture from wild aurochs (Park et al., 2015; Upadhyay et al., 2017). This observation highlights the genetic distinctiveness of the Kerry population and has major implications for conservation and management of the breed.

For the present study, and within a genetic conservation framework, we performed high-resolution comparative population genomics analyses of Kerry cattle and a range of British and European cattle breeds. These analyses encompassed phylogenetic network reconstruction, evaluation of genetic structure and inbreeding, modelling of historical effective population sizes and functional analyses of artificial and natural selection across the Kerry genome.

## 2. Materials and Methods

### 2.1. Kerry Cattle Population DNA Sampling in 1991/92 and 2011/12

Two different population samples from the Irish Kerry cattle breed were used for this study (**Figure 1**). The first population sample consisted of peripheral blood and semen straw genomic DNA collected and purified from 36 male and female Kerry cattle in 1991/92, which are a subset of the Kerry cattle population sample (*n* = 40) we have previously described and used for microsatellite-based population genetics analyses (MacHugh et al., 1997; MacHugh et al., 1998). Pedigree records and owners were consulted to ensure there was the minimum degree of genetic relatedness among the animals sampled. This Kerry population sample group is coded as KY92.

**FIGURE 1.**
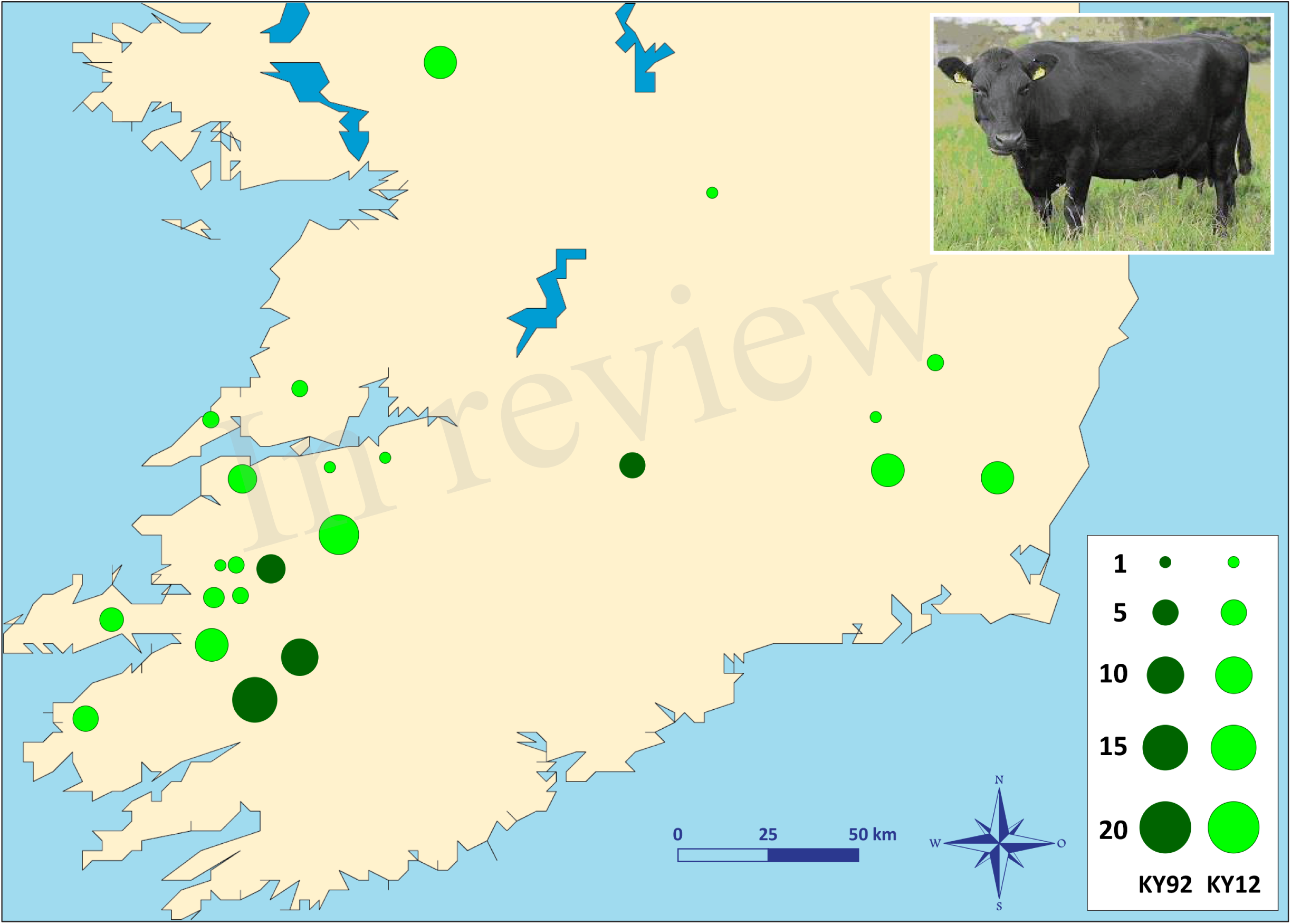
Photograph of a Kerry cow and locations of Kerry cattle herd DNA sampling in southern Ireland. The area of each circle corresponds to the size of each population sample. Dark green = animals sampled during 1991/92 (KY92); light green = animals sampled during 2011/12 (KY12). Kerry cow image is copyright of the Kerry Cattle Society Ltd.

The second Kerry cattle population sample was collected in 2011/12 from 19 different herds located across southern and western Ireland. Performagene (PG-100) nasal swab DNA collection kits were used for biological sample collection (DNA Genotek Inc., Ottawa, Canada). Nasal swab DNA samples were collected from a total of 75 male and female Kerry cattle and owners were consulted to ensure there was the minimum degree of genetic relatedness among the animals sampled. This Kerry population sample group is coded as KY12. Genomic DNA was purified from nasal swabs using the Promega Maxwell^®^ 16 DNA automated DNA extraction platform (Promega UK, Southampton, United Kingdom) and established methods developed at the UCD Animal Genomics Laboratory (Magee et al., 2010; Magee et al., 2011).

### 2.2 SNP Genotyping and Assembly of Comparative SNP Data Sets

Illumina^®^ Bovine SNP50 BeadChip (Matukumalli et al., 2009) genotyping on all 111 Kerry genomic DNA samples (KY92 and KY12 sample panels plus nine blinded sample duplicates for quality control purposes) was performed by Weatherbys Scientific (Co. Kildare, Ireland).

For comparative population genomics analyses, equivalent SNP data for a range of other breeds were obtained from previously published work (Decker et al., 2009; Flori et al., 2009; Gibbs et al., 2009; Matukumalli et al., 2009; Gautier et al., 2010; Park et al., 2015). The breed SNP data were split into two discrete composite data sets: a European breed SNP data set (EU) and a SNP data set for a subset of European breeds originating from Britain and Ireland (BI). A population sample of West African N’Dama *B. taurus* cattle from Guinea (NDAM) was also used as an outgroup for the phylogenetic analyses. **Table 1** provides detailed biogeographical information on the cattle breed samples used for the present study.

**Table 1:** Cattle breed/population samples used for the present study

| Breed/population | Code | Data set | Breed purpose | Country of origin | Source |
| --- | --- | --- | --- | --- | --- |
| Angus | ANGU | BI/EU | Beef | Scotland | 1, 2 |
| Belted Galloway | BGAL | BI | Beef | Scotland | 2 |
| British Shorthorn | BSHN | BI/EU | Dual purpose | England | 2 |
| Brown Swiss | BRSW | EU | Dairy | Switzerland | 1, 2, 3 |
| Charolais | CHAR | EU | Beef | France | 1, 2, 3 |
| Devon | DEVN | BI | Beef | England | 2 |
| Dexter | DXTR | BI | Dual purpose | Ireland | 2 |
| English Longhorn | ELHN | BI | Beef | England | 2 |
| Finnish Ayrshire | FAYR | BI/EU | Dairy | Scotland/Finland | 2 |
| Galloway | GALL | BI | Beef | Scotland | 2 |
| Gelbvieh | GELB | EU | Dual purpose | Germany | 2, 4 |
| Guernsey | GNSY | BI/EU | Dairy | Channel Islands | 1, 2 |
| Hereford | HRFD | BI/EU | Beef | England | 1, 2 |
| Holstein | HOLS | EU | Dairy | The Netherlands | 1, 2, 5 |
| Jersey | JRSY | BI/EU | Dairy | Channel Islands | 1, 2, 3 |
| Kerry sampled 1991/92 | KY92 | BI/EU | Dairy | Ireland | Current |
| Kerry sampled 2011/12 | KY12 | BI/EU | Dairy | Ireland | Current |
| Limousin | LIMS | EU | Beef/draft | France | 1, 2 |
| Lincoln Red | LNCR | BI | Beef | England | 2 |
| Montbeliarde | MONT | EU | Dairy | France | 2, 5 |
| N'Dama | NDAM | --- | Dual purpose | Guinea (West Africa) | 1 |
| Norwegian Red | NRED | EU | Dairy | Norway | 1 |
| Piedmontese | PDMT | EU | Dual purpose | Italy | 1, 2 |
| Red Angus | RANG | BI/EU | Beef | Scotland | 1 |
| Red Poll | REDP | BI | Beef | England | 2 |
| Romagnola | ROMG | EU | Beef/draft | Italy | 1 |
| Scottish Highland | SCHL | BI | Beef | Scotland | 2 |
| Simmental | SIMM | EU | Dual purpose/draft | Switzerland | 2, 4 |
| South Devon | SDEV | BI | Beef | England | 2 |
| Sussex | SUSX | BI | Beef/draft | England | 2 |
| Welsh Black | WBLK | BI | Dual purpose | Wales | 2 |
| White Park | WHPK | BI | Dual purpose/draft | England | 2 |

<sup>1</sup> Gibbs et al. (2009); <sup>2</sup> Decker et al. (2009); <sup>3</sup> Gautier et al. (2010); <sup>4</sup> Matukumalli et al. (2009); <sup>5</sup> Flori et al. (2009)

### 2.3. Sample Removal and Quality Control and Filtering of SNPs

Pedigree records and genomic relationship were used to identify animals from the KY92 and KY12 population samples that were either parent-offspring pairs or full-sibs. One of the two animals in each pair was then randomly removed to generate the working SNP data set. Following this procedure, quality control and filtering based on recorded SNP genotypes was performed as detailed below for the EU and BI data sets.

Prior to quality control and filtering there were 54,057 SNPs in the EU data set (608 animals, including KY92 and KY12) and in the BI data set (354 animals, including KY92 and KY12). SNP quality filtering was performed using PLINK version 1.07 (Purcell et al., 2007), such that individual SNPs with more than 10% missing data and a minor allele frequency (MAF) of ≤ 0.01 were removed from both data sets. X and Y chromosome SNPs were removed and individual animal samples with a SNP call rate less than 90% were also removed from each of the two data sets.

SNP quality control and filtering were performed across breeds/populations (by data set) for construction of phylogenies and ancestor graphs, multivariate analysis, investigation of population structure and detection of signatures of selection. For intrapopulation analyses of effective population size (*N*_e_) and genomic inbreeding, all SNPs genotyped (54,057) were filtered within breeds/populations as detailed above. However, an additional filtering procedure was used to remove SNPs deviating from Hardy-Weinberg equilibrium (HWE) with a *P* value threshold of < 0.0001. Also, for the *N*_e_ analysis, a more stringent MAF threshold of 0.05 was used.

Using the filtered genome-wide SNP data, PLINK v1.07 was also used to generate identity-by-state (IBS) values between all pairs of Kerry cattle (KY92 and KY12), including the nine blinded sample duplicates for quality control purposes.

### 2.4. Construction of Phylogenetic Trees and Ancestry Graphs

Maximum likelihood (ML) phylogenetic trees with ancestry graphs were generated for the EU and BI data sets using the TreeMix (version 1.12) software package (Pickrell and Pritchard, 2012). The West African *B. taurus* NDAM breed sample (*n* = 22) was used as an outgroup. TreeMix was run without using SNP blocks (as described in the TreeMix software documentation) and ML phylogenetic trees were generated with no migration edges (*m* = 0) up to ten migration edges (*m* = 10).

### 2.5. Population Differentiation and Genetic Structure

To visualise the main axes of genomic variation among cattle breeds and individual animals, multivariate principal component analysis (PCA) was performed for the composite EU and BI SNP data sets using SMARTPCA from the EIGENSOFT package (version 4.2) with default settings (Patterson et al., 2006).

To further investigate genetic structure and admixture history for Kerry cattle and other breeds the fastSTRUCTURE software package (Raj et al., 2014) was used to analyse the EU and BI data sets for a range of *K* possible ancestral populations (*K* = 2‒15). For the present study, the simple prior approach described by Raj and colleagues (2014) was used, which is sufficient for modelling population/breed divergence. To identify the ‘true’ *K* value for the number of ancestral populations, a series of fastSTRUCTURE runs with pre-defined *K* values were examined using the *chooseK.py* script (Raj et al., 2014). Outputs from the fastSTRUCTURE analyses were visualised using the DISTRUCT software program (Rosenberg et al., 2002) using standard parameters.

### 2.6. Modelling Current and Historical Effective Population Size (*N*_e_)

Current and historical *N*_e_ trends were modelled with genome-wide SNP linkage disequilibrium (LD) data for the KY92 and KY12 populations plus a selection of BI and EU breeds using the SNeP software tool as described by Barbato and colleagues (Barbato et al., 2015). This method facilitates estimation of historical *N*_e_ values from SNP linkage disequilibrium (LD) data using the following equation (Corbin et al., 2012):

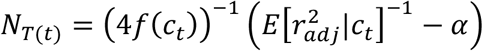

where *N*_*T*_ is the effective population size *t* generations ago calculated as *t* = (2*f*(*c*_*t*_))^−1^ (Hayes et al., 2003), *c*_*t*_ is the recombination rate defined for a specific physical distance between SNP markers, 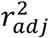 is the LD value adjusted for sample size and *α* := {1, 2, 2.2} is a correction for the occurrence of mutation (Barbato et al., 2015). In addition, the SNeP program option for unphased SNP data was used for the analyses described here.

### 2.7. Evaluation of Genomic Inbreeding and Runs of Homozygosity (ROH)

Individual animal genomic inbreeding was evaluated as genome-wide autozygosity estimated from SNP data using runs of homozygosity (ROH) and the *F*_ROH_ statistic introduced by McQuillan and colleagues (2008). The *F*_ROH_ statistic was calculated as the ratio of the total length of defined runs of homozygosity (*L*_ROH_) to the total length of the autosomal genome covered by SNPs:

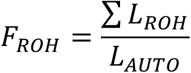

PLINK v1.07 was used to define runs of homozygosity (ROH) using a sliding window approach and procedures modified from previous recommendations for Illumina^®^ Bovine SNP50 BeadChip and similar SNP data sets (Purfield et al., 2012; Purfield et al., 2017). The criteria for defining individual ROH were set such that the ROH was required to be at least 500 kb in length, with a minimum density of one SNP per 120 kb and that there was a gap of at least 1,000 kb between each ROH. A sliding window of 50 SNPs was incrementally advanced one SNP at a time along the genome; each discrete window could contain a maximum of one heterozygous SNP and no more than two SNPs with missing genotypes. Following Purfield and colleagues all filtered genomic SNPs, including those located in centromeric regions, were used to estimate *F*_ROH_ values for individual animals.

### 2.8. Genome-wide Detection of Signatures of Selection and Functional Enrichment Analysis

In the absence of hard selective sweeps, single selection tests using high-density SNP data do not perform well in detecting signatures of selection from individual livestock breeds (Kemper et al., 2014). Therefore, for the present study, genomic signatures of selection were identified using the composite selection signal (CSS) method introduced by Randhawa and colleagues (Randhawa et al., 2014). The CSS method has been shown to be a robust and sensitive approach for detecting genomic signatures of selection underlying microevolution of complex traits in livestock (Randhawa et al., 2015). The CSS is a weighted index of signatures of selection from multiple estimates; it is a nonparametric procedure that uses fractional ranks of constituent tests and does not depend on assumptions about the distributions of individual test results.

As described in detail by Randhawa *et al.* (2014), the CSS method can be used to combine the fixation index (*F*_ST_), the change in selected allele frequency (Δ*SAF*) and the cross-population extended haplotype homozygosity (*XP-EHH*) tests into one composite statistic for each SNP in a population genomics data set. For the present study, we used 36,621 genome-wide SNPs genotyped in 98 individual Kerry cattle samples (from both the KY92 and KY12 populations) and a sample of 102 randomly selected cattle (six random cattle from each breed of the EU data set). To mitigate against false positives, genomic selection signatures were only considered significant if at least one SNP from the set of the top 0.1% genome-wide CSS scores was flanked by at least five SNPs from the set of the top 1% CSS scores.

The Ensembl BioMart data mining resource (Smedley et al., 2015) was used to identify genes within ± 1.0 Mb of each selection peak (Ensembl release 90, August 2017). Following this, Ingenuity^®^ Pathway Analysis (IPA^®^: Qiagen, Redwood City, CA, USA; release date June 2017) was used to perform an overrepresentation enrichment analysis (ORA) with this gene set to identify canonical pathways and functional processes of biological importance. The total gene content of Ensembl release 90 version of the UMD3.1 bovine genome assembly (Zimin et al., 2009) was used as the most appropriate reference gene set for these analyses (Timmons et al., 2015).

## 3. Results and Discussion

### 3.1. Sample Removal and SNP Filtering and Quality Control

Inspection of the pedigree records and the genomic relationship matrix (based on identity-by-state [IBS] of SNP genotypes) identified 10 animals from the KY12 population that were members of a parent-offspring or full-sib pair (sample codes: KY12_01, KY12_05, KY12_13, KY12_14, KY12_17, KY12_18, KY12_19, KY12_46, KY12_55, KY12_67). One sample from each of these pairs was randomly removed. Thereafter, general SNP quality control and filtering led to additional samples being excluded (KY12_26, KY12_28 and KY12_54), giving a total filtered KY12 population sample of 62 animals for downstream population genomics analyses.

After SNP quality control and filtering across the two composite data sets (EU and BI), there were 36,621 autosomal SNPs from 605 individual animals in the EU data set and there were 37,395 autosomal SNPs from 351 animals in the BI data set. When the West African NDAM breed sample (*n* = 22) was included for the ML phylogenetic tree and ancestry graph analyses, the number of SNPs used was 36,000 from 627 animals for the EU data set and 37,490 from 373 animals for the BI data set. The final numbers of SNPs used for individual breed/population analyses of *N*_e_ and genomic inbreeding after all quality control and filtering (including additional filtering for deviations from HWE) are shown in **Table 2**.

**Table 2:**
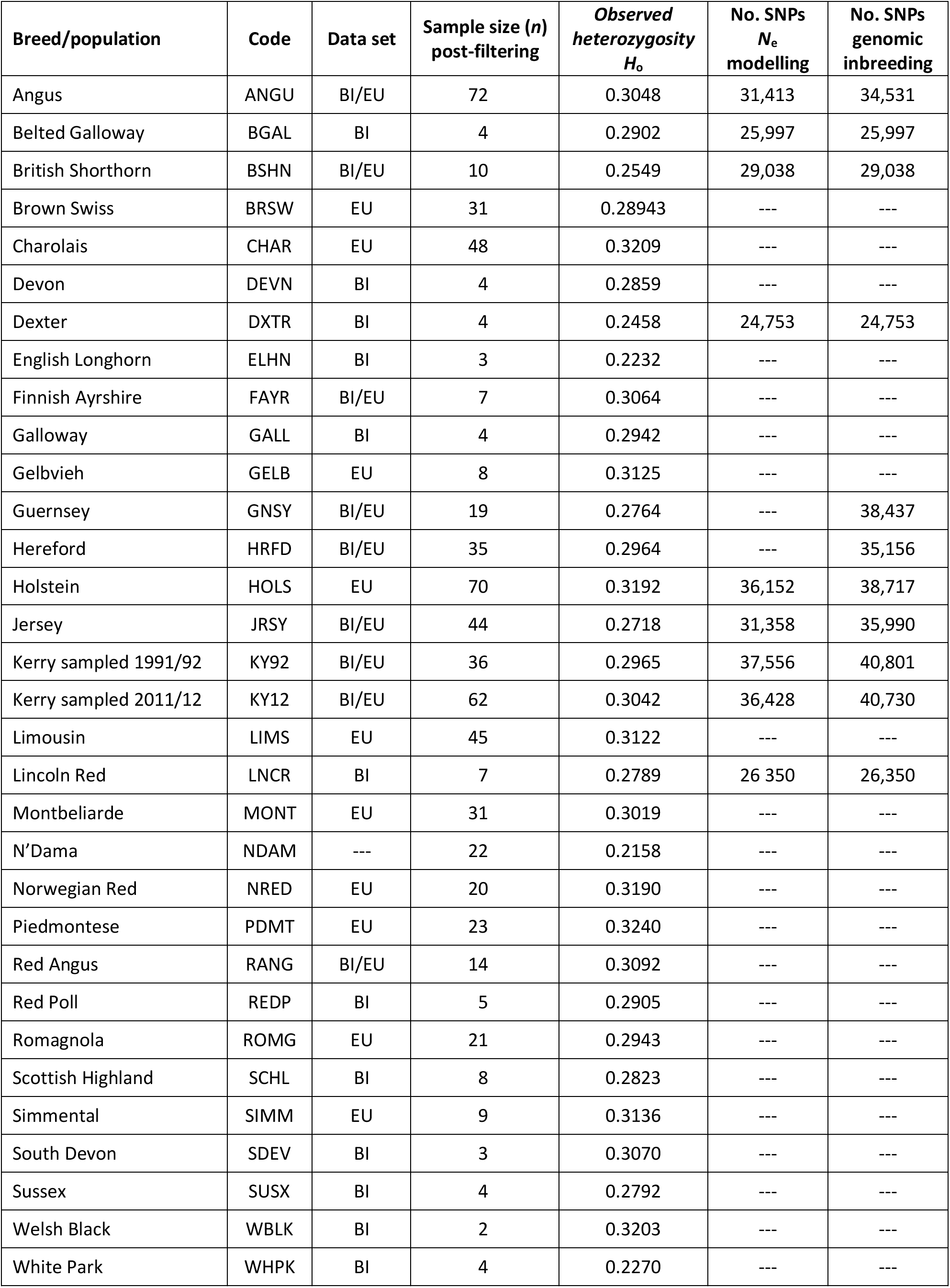
Breed/population sample size, observed heterozygosity and SNP filtering information

| Breed/population | Code | Data set | Sample size (*n*)<br>post-filtering | Observed<br>heterozygosity<br>*H<sub>o</sub>* | No. SNPs<br>*N<sub>e</sub>*<br>modelling | No. SNPs<br>genomic<br>inbreeding |
| --- | --- | --- | --- | --- | --- | --- |
| Angus | ANGU | BI/EU | 72 | 0.3048 | 31,413 | 34,531 |
| Belted Galloway | BGAL | BI | 4 | 0.2902 | 25,997 | 25,997 |
| British Shorthorn | BSHN | BI/EU | 10 | 0.2549 | 29,038 | 29,038 |
| Brown Swiss | BRSW | EU | 31 | 0.28943 | --- | --- |
| Charolais | CHAR | EU | 48 | 0.3209 | --- | --- |
| Devon | DEVN | BI | 4 | 0.2859 | --- | --- |
| Dexter | DXTR | BI | 4 | 0.2458 | 24,753 | 24,753 |
| English Longhorn | ELHN | BI | 3 | 0.2232 | --- | --- |
| Finnish Ayrshire | FAYR | BI/EU | 7 | 0.3064 | --- | --- |
| Galloway | GALL | BI | 4 | 0.2942 | --- | --- |
| Gelbvieh | GELB | EU | 8 | 0.3125 | --- | --- |
| Guernsey | GNSY | BI/EU | 19 | 0.2764 | --- | 38,437 |
| Hereford | HRFD | BI/EU | 35 | 0.2964 | --- | 35,156 |
| Holstein | HOLS | EU | 70 | 0.3192 | 36,152 | 38,717 |
| Jersey | JRSY | BI/EU | 44 | 0.2718 | 31,358 | 35,990 |
| Kerry sampled 1991/92 | KY92 | BI/EU | 36 | 0.2965 | 37,556 | 40,801 |
| Kerry sampled 2011/12 | KY12 | BI/EU | 62 | 0.3042 | 36,428 | 40,730 |
| Limousin | LIMS | EU | 45 | 0.3122 | --- | --- |
| Lincoln Red | LNCR | BI | 7 | 0.2789 | 26 350 | 26,350 |
| Montbeliarde | MONT | EU | 31 | 0.3019 | --- | --- |
| N'Dama | NDAM | --- | 22 | 0.2158 | --- | --- |
| Norwegian Red | NRED | EU | 20 | 0.3190 | --- | --- |
| Piedmontese | PDMT | EU | 23 | 0.3240 | --- | --- |
| Red Angus | RANG | BI/EU | 14 | 0.3092 | --- | --- |
| Red Poll | REDP | BI | 5 | 0.2905 | --- | --- |
| Romagnola | ROMG | EU | 21 | 0.2943 | --- | --- |
| Scottish Highland | SCHL | BI | 8 | 0.2823 | --- | --- |
| Simmental | SIMM | EU | 9 | 0.3136 | --- | --- |
| South Devon | SDEV | BI | 3 | 0.3070 | --- | --- |
| Sussex | SUSX | BI | 4 | 0.2792 | --- | --- |
| Welsh Black | WBLK | BI | 2 | 0.3203 | --- | --- |
| White Park | WHPK | BI | 4 | 0.2270 | --- | --- |

All data sets, including EU and BI composite data sets and individual breed/population data sets had total SNP call rates of > 99%. The IBS values estimated for Kerry cattle (KY92 and KY12) from filtered genome-wide SNP data are reported in Supplementary Table 1 and described further in **Section 3.7**.

### Observed heterozygosity (*H*_o_) estimated from genome-wide SNP data

**Table 2** provides genome-wide *H*_o_ values for each of the breeds/populations used for the present study. The lowest genome-wide *H*_o_ value was observed for the West Africa NDAM *B. taurus* breed, which is likely a consequence of ascertainment bias introduced by a focus on polymorphic SNPs in European *B. taurus* during design of the Illumina^®^ Bovine SNP50 BeadChip (Matukumalli et al., 2009).

Generally, as shown in **Table 2** for the EU and BI breeds and populations, local landrace or heritage breeds display lower *H*_o_ values compared to more widespread production breeds such as the Simmental (SIMM), Holstein (HOLS) or Charolais (CHAR) breeds. In addition, as might be expected, production breeds originally derived from minor island populations (Jersey [JRSY] and Guernsey [GNSY]) also exhibit relatively low *H*_o_ values. In the context of genetic conservation it is therefore encouraging that the KY92 and KY12 population samples display intermediate *H*_o_ values that are at the upper end of the range observed for the heritage breeds.

### 3.3. Maximum Likelihood Phylogenetic Ancestry Graphs using Genome-wide SNP Data

To examine microevolutionary patterns of genetic differentiation and gene flow among cattle breeds and populations, ML phylogenetic ancestry graphs were generated using TreeMix. For the EU data set, the ML tree topology was consistent for all values of *m*, with the exception of *m* = 2 migration edges, where the Hereford breed (HRFD) was observed to group with the HOLS breed. The ML tree generated with *m* = 5 is shown in **Figure 2**, which highlights the genetic similarity of the Northern European breeds (British, Irish and Scandinavian). As expected the two Kerry population samples (KY92 and KY12) are genetically very similar and emerge on the same branch as the HRFD breed. It is also noteworthy that there is a high-weight migration edge between the British Shorthorn breed (BSHN) and the root of the two Kerry population samples, supporting the hypothesis of historical gene flow from the British Shorthorn breed into the ancestral population of modern Kerry cattle (Curran, 1990).

**FIGURE 2.**
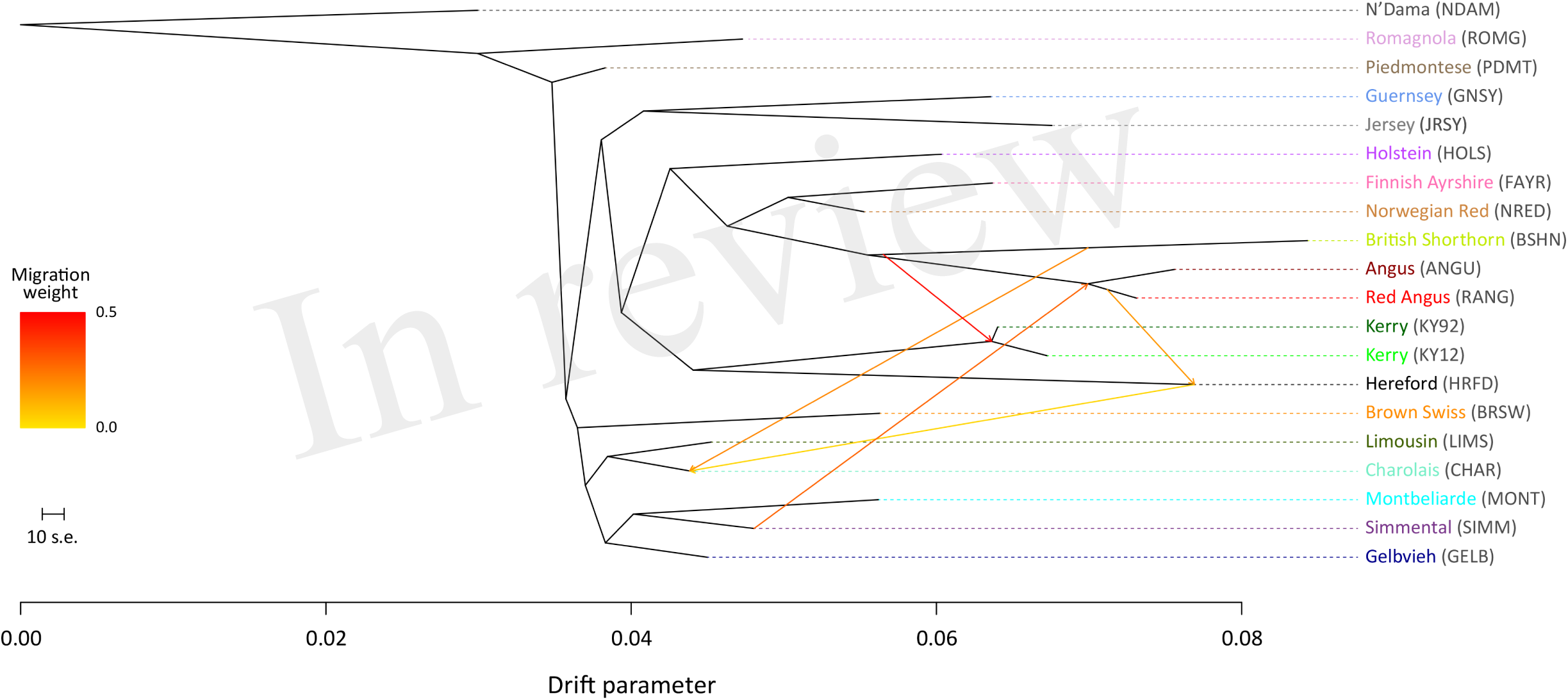
Maximum likelihood (ML) phylogenetic tree network graph with five migration edges (*m* = 5) generated for genome-wide SNP data (36,000 autosomal SNPs) from European cattle breeds (EU data set). The West African taurine N’Dama breed sampled in Guinea is included as a population outgroup. Coloured lines and arrows show migration edges that model gene flow between lineages with different migration weights represented by the colour gradient.

For the ML trees generated using the BI data set, breed/population differentiation was less apparent, possibly due to similar biogeographical origins for these breeds and/or smaller sample sizes for some of the populations sampled. **Figure 3** shows the ML tree generated with *m* = 5 for the BI data set. For *m* = 5, all migration edges stem from the BSHN/Lincoln Red (LNCR) branch, including a medium-weight migration edge connecting to the Kerry cattle branch. These results support the hypothesis that there was significant gene flow during the 18^th^ and 19^th^ centuries from British Shorthorn cattle into the ancestral populations for a range of modern British and Irish cattle breeds (Grobet et al., 1998; Felius et al., 2011; Felius et al., 2015).

**FIGURE 3.**
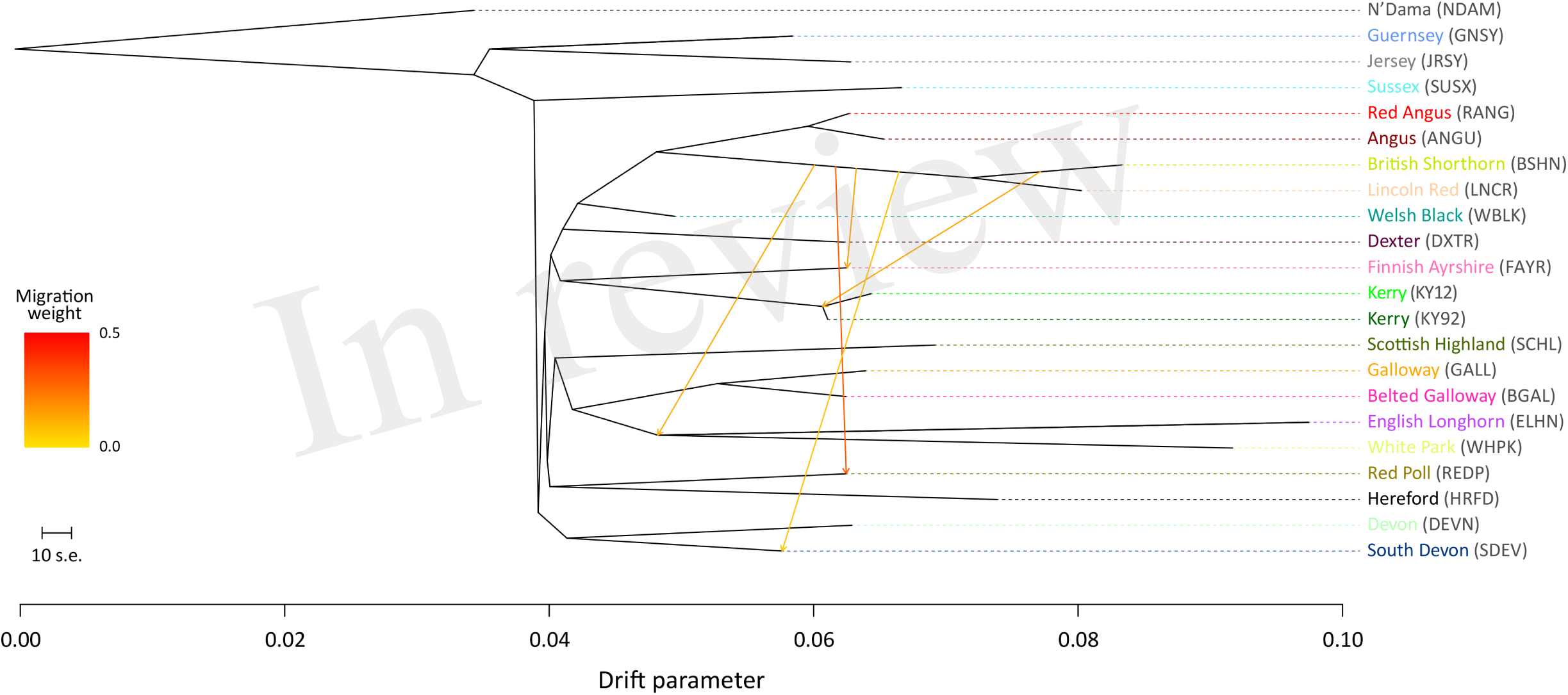
Maximum likelihood (ML) phylogenetic tree network graph with five migration edges (*m* = 5) generated for genome-wide SNP data (37,490 autosomal SNPs) from cattle breeds of British and Irish origin (BI data set). The West African taurine N’Dama breed sampled in Guinea is included as a population outgroup. Coloured lines and arrows show migration edges that model gene flow between lineages with different migration weights represented by the colour gradient.

### 3.4. Multivariate Principal Component Analysis of Genome-wide SNP Data

To investigate inter-and intra-population genomic diversity and genetic relationship among individual animals from multiple cattle breeds and populations, PCA was performed using genome-wide SNP data. Principal component plots of the first (PC1) and second (PC2) principal components are shown in **Figures 4** and **5** for the EU and BI data sets, respectively.

In **Figure 4**, for the EU data set, PC1 and PC2 account for 18.2% and 16.8% of the total variation for PC1–10, respectively. The PC1 plot axis differentiates the British Angus (ANGU), Red Angus (RANG) and BSHN and Irish KY92 and KY12 populations from the rest of the European breeds, including the British HRFD and GNSY and JRSY Channel Islands breeds. In addition, the ANGU and RANG and the Kerry (KY92 and KY12) emerge at the opposite extremes of the PC2 plot axis, highlighting the genetic distinctiveness of the Kerry breed and supporting their status as an important cattle genetic resource that should be prioritised for conservation.

**FIGURE 4.**
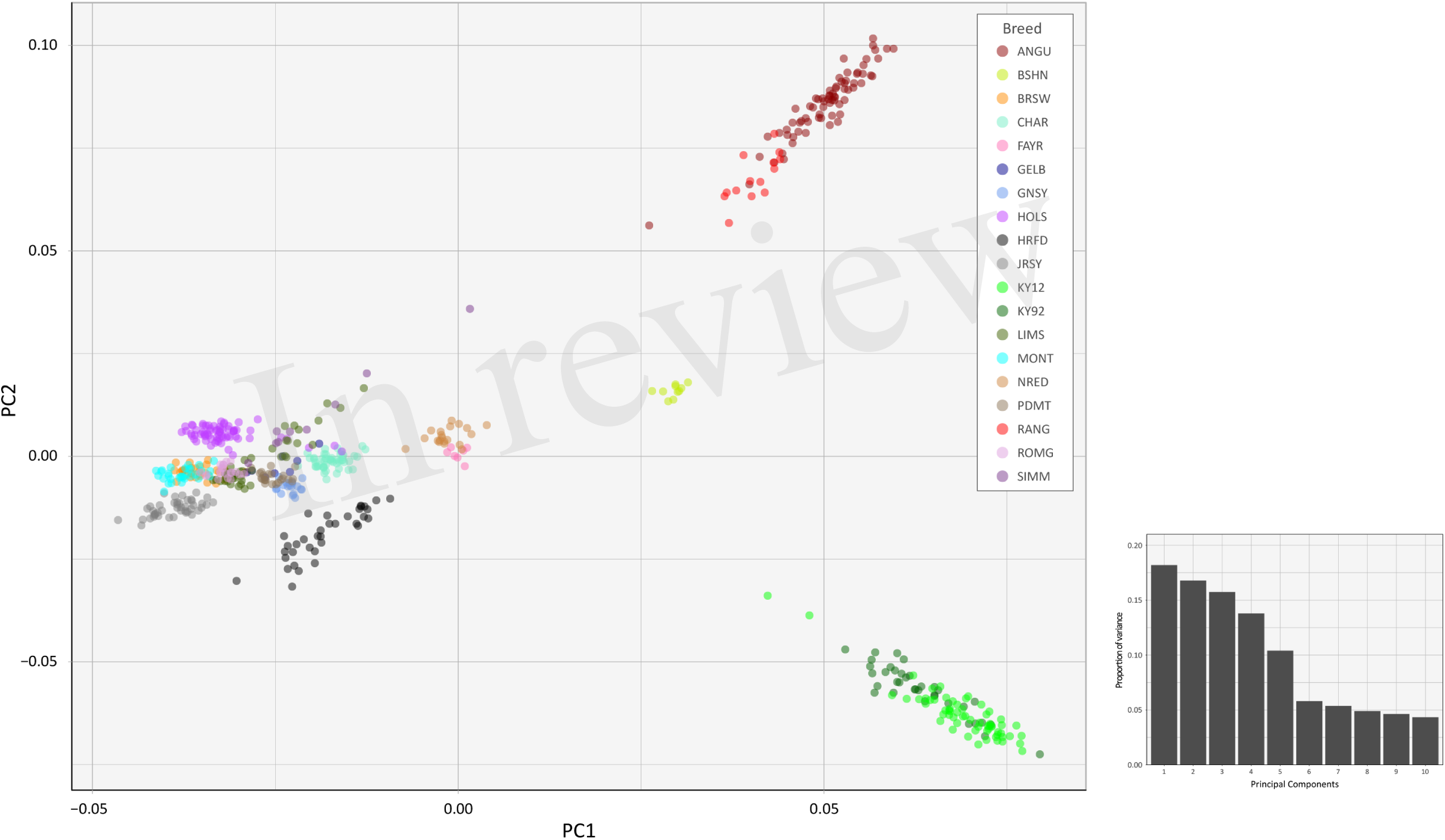
Principal component analysis plot constructed for PC1 and PC2 from genome-wide SNP data (36,621 autosomal SNPs) for the EU data set of 605 individual animals. The smaller histogram plot shows the relative variance contributions for the first 10 PCs.

In **Figure 5**, for the BI data set, PC1 and PC2 account for 23.7% and 22.7% of the total variation for PC1–10, respectively. The PC1 plot axis recapitulates PC2 in **Figure 4** and differentiates the Kerry (KY92 and KY12) from the ANGU and RANG breeds with the other British breeds emerging between these two extremes. This result reiterates the genetic distinctiveness of the Kerry cattle breed in comparison to a wide range of British production and heritage landrace cattle breeds, again emphasising the need for genetic conservation.

**FIGURE 5.**
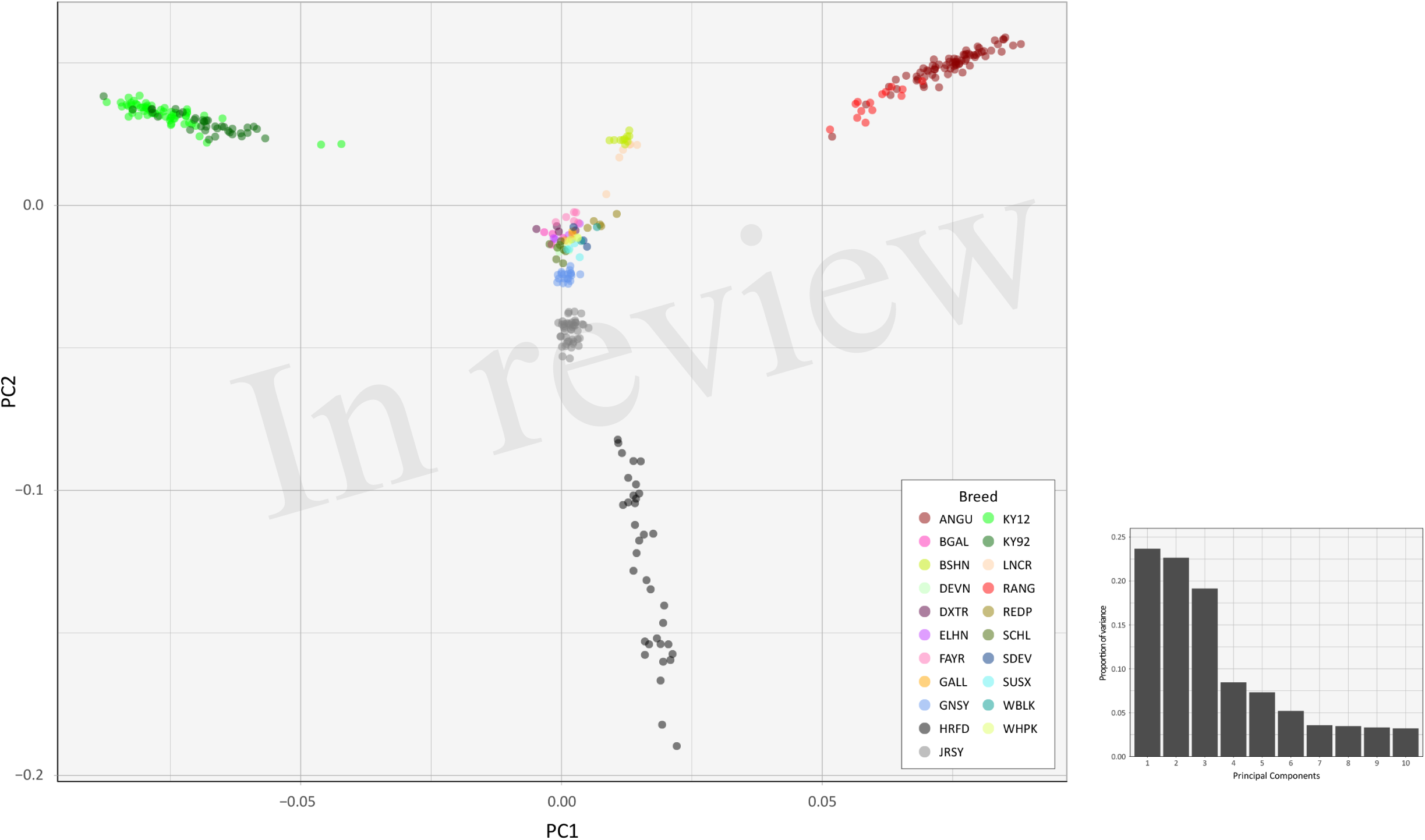
Principal component analysis plot constructed for PC1 and PC2 from genome-wide SNP data (37,395 autosomal SNPs) for the BI data set of 351 individual animals. The smaller histogram plot shows the relative variance contributions for the first 10 PCs.

The PC2 plot axis in **Figure 5** differentiates the HRFD breed from the other British and Irish breeds and reveals substantial genetic diversity among individual HRFD animals. However, in this context, it is important to note that the pattern of genetic diversity revealed here for the HRFD population sample may be due to ascertainment bias as a consequence of the strategy used to design the Illumina^®^ Bovine SNP50 BeadChip. In this regard, many of the SNPs that constitute this first-generation SNP array were identified from heterozygous positions in the inbred Hereford female (L1 Dominette 01449) bovine genome assembly or through comparisons of random shotgun reads from six diverse cattle breeds that were aligned directly to the same Hereford genome assembly (Matukumalli et al., 2009). This approach to SNP array design will inevitably lead to elevated intrabreed genomic variation using the Illumina^®^ Bovine SNP50 BeadChip with Hereford cattle (Meuwissen, 2009) and accounts for the dispersed pattern of individual HRFD samples in **Figure 5**.

Examination of **Figures 4** and **5** indicates that two of the KY12 animals sampled may exhibit a genetic signature of ancestral crossbreeding with another cattle population, which, anecdotally, is likely to have been due to crossbreeding with Angus cattle. Therefore, another PCA plot was generated (Supplementary Figure 1) that shows PC1 and PC2 for individual animals from the KY92, KY12, ANGU and RANG population samples. The two animals exhibiting a genetic signature of possible ancestral crossbreeding (KY12_06 and KY12_58) are indicated on Supplementary Figure 1. Under the assumption of crossbreeding with ANGU cattle, the positions of KY12_06 and KY12_58 on this PC plot would suggest that these animals are third generation (F_3_) or greater backcrosses. Notwithstanding the KY12_06 and KY12_58 data points, the genetic similarity among all Kerry cattle sampled is evident by comparison of the tight KY92 and KY12 sample cluster to the dispersion of the ANGU and RANG samples on the PCA plot in Supplementary Figure 1.

### 3.5. Analysis of Genetic Structure using Genome-wide SNP data

The results of the fastSTRUCTURE analyses using the EU and BI data sets are shown in **Figure 6** and **Figure 7**, respectively. For both analyses, the Kerry cattle (KY92 and KY12) cluster as a single group at *K* = 2 and are differentiated from all other European or British and Irish cattle breeds. The other breed group that is clearly differentiated at *K* = 2 in **Figure 7** is the cluster composed of the ANGU and RANG breeds. These results mirror the pattern shown for PC1 in **Figure 5**, and again emphasise the genetic distinctiveness of Kerry cattle compared to other European production and landrace heritage breeds. Using the *chooseK.py* script the ‘true’ number of clusters corresponding to the likely number of ancestral populations was estimated to be between 12 and 14 for the EU data set and either 7 or 8 for the BI data set.

**FIGURE 6.**
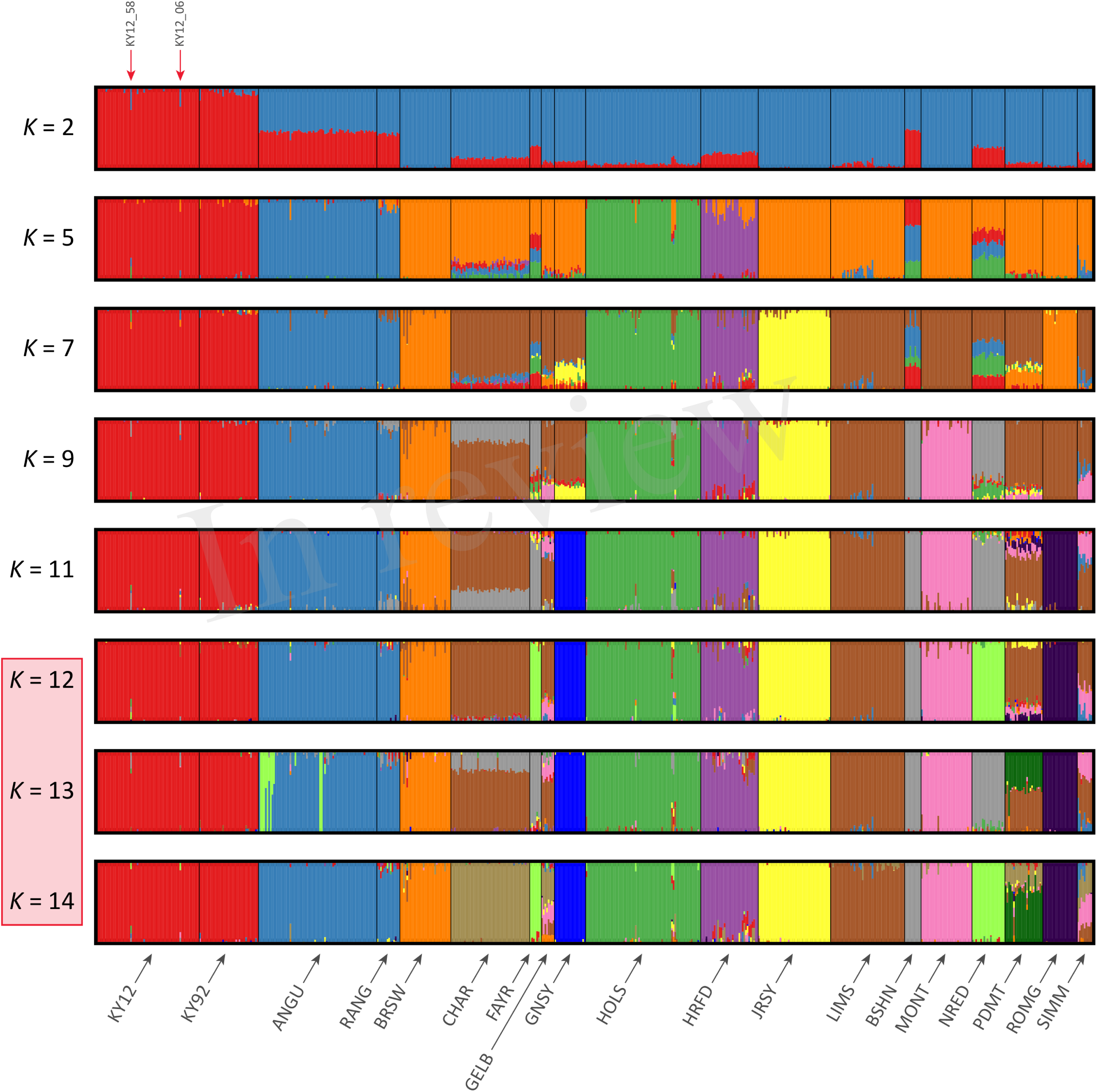
Hierarchical clustering of individual animals using genome-wide SNP data (36,621 autosomal SNPs) for the EU data set of 605 individual animals. Results are shown for modelled ancestral populations *K* = 2 to 14. The cluster numbers corresponding to the likely number of ancestral populations are highlighted with a light red overlay and the two outlier Kerry samples (KY12_06 and KY12_58) are indicated with red arrows.

**FIGURE 7.**
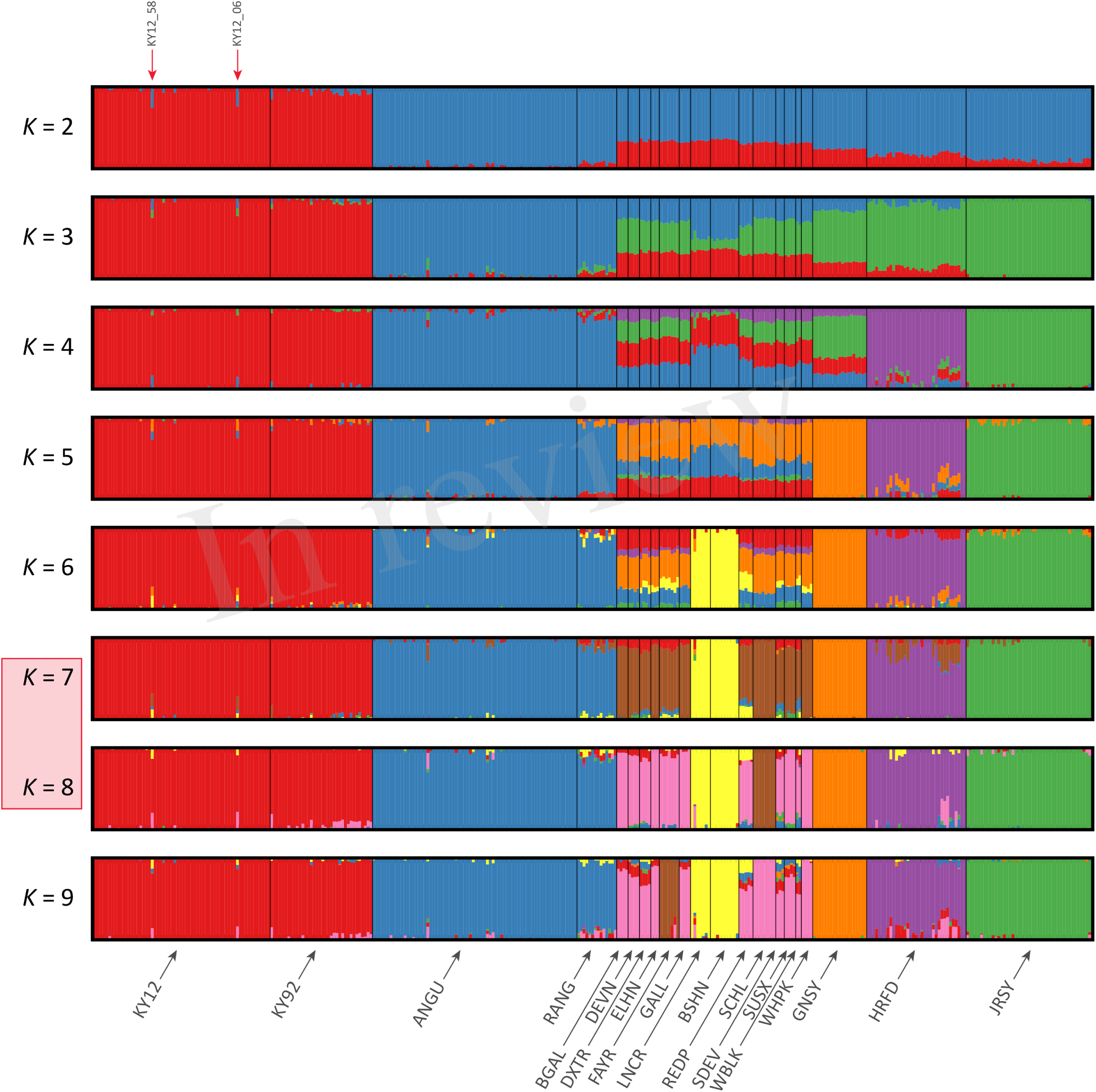
Hierarchical clustering of individual animals using genome-wide SNP data (37,395 autosomal SNPs) for the BI data set of 351 individual animals. Results are shown for modelled ancestral populations *K* = 2 to 9. The cluster numbers corresponding to the likely number of ancestral populations are highlighted with a light red overlay and the two outlier Kerry samples (KY12_06 and KY12_58) are indicated with red arrows.

For both data sets, animals from the KY12 population sample appear to be more genetically homogenous compared to the KY92 population sample. This observation may be a consequence of increasing use, since the early 1990s, of small numbers of artificial insemination (AI) Kerry sires. It is also noteworthy that the two individual animals detected with a substantial signature of putative historical crossbreeding (KY12_06 and KY12_58) show marked patterns of population admixture in the fastSTRUCTURE results, which are indicated by red arrows in **Figure 6** and **Figure 7**.

### 3.6. Modelling Historical Effective Population Size (*N*_e_) using Genome-wide SNP Data

The results from modelling historical *N*_e_ in a selection of production and heritage cattle breeds and populations (KY92, KY12, DXTR, BSHN, BGAL, LNCR, ANGU, JRSY and HOLS) are provided in Supplementary Table 2 and visualised in **Figure 8**. The ‘demographic fingerprints’ (Barbato et al., 2015) of the two Kerry populations shown in **Figure 8** and tabulated in Supplementary Table 2 are more similar to those of the production breeds with large census populations (BSHN, ANGU, JRSY, HOLS) than the other heritage breeds with relatively small census population sizes (DXTR, BGAL, LNCR). The KY92, KY12, BSHN, ANGU, JRSY and HOLS populations show a declining trend from historical *N*_e_ peaks between 1,500 and 2,000 more than 900 generations ago to *N*_e_ values estimated to be less than 200 within the last 20 generations. One the other hand, the DXTR, BSHN, BGAL and LNCR populations display a more severe decline from historical *N*_e_ peaks between 2,500 and 4,000 more than 900 generations ago to *N*_e_ values estimated to be less than 150 within the last 20 generations.

**FIGURE 8.**
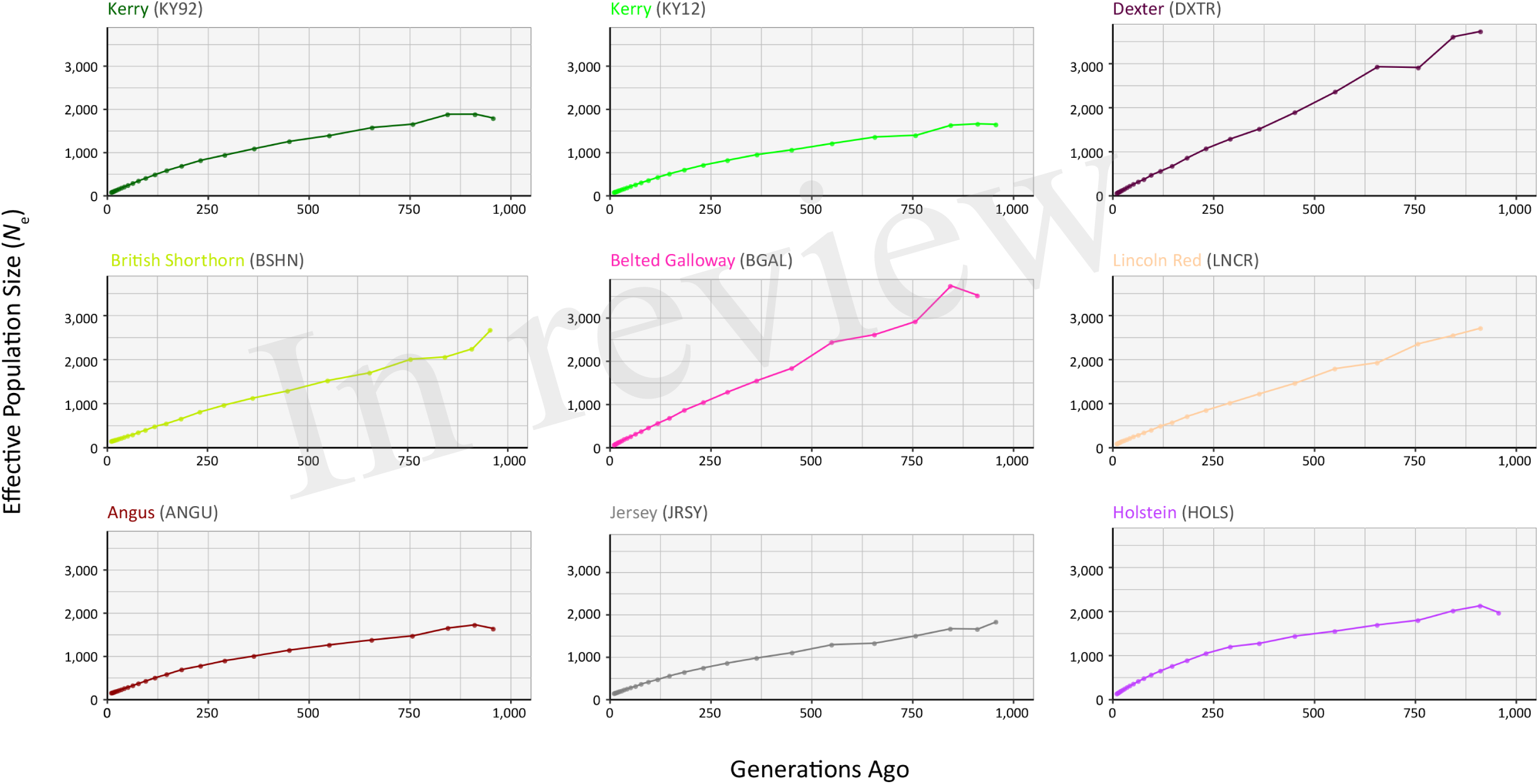
Genetic effective population size (*N*_e_) trends modelled using genome-wide SNP data. Results for the KY92 and KY12 populations are shown with seven comparator heritage and production cattle breeds.

It is important to keep in mind these *N*_e_ trends may be partly a consequence of the relatively small sample sizes for the DXTR, BGAL and LNCR breeds (see **Table 2**), coupled with different histories of migration, gene flow and, in particular, strong artificial selection in the production cattle populations. Notwithstanding these caveats, the most recent modelled *N*_e_ values for the KY92 and KY12 population samples are 89 and 88, respectively. These values are *N*_e_ estimates for 12 generations in the past and assuming a generation interval of between 4 to 6 years, which is based on a pedigree estimate from a similar heritage cattle population of 5.66 (Meszaros et al., 2015), this corresponds to between 48 and 72 years before 2012 (for the KY12 population). This is approximately the period between 1940 and 1965, which is during the time that the Kerry breed started to decline precipitously in census population size and also *N*_e_ estimated from herd book data (O'hUigín and Cunningham, 1990; Food and Agriculture Organization, 2017).

From a conservation perspective, livestock populations generally exhibit *N*_e_ values relative to total census population sizes (*N*_c_) that are substantially lower than seen in comparable wild mammal populations (Hall, 2016). Also, estimation of *N*_e_ using methods such as SNeP that leverage genome-wide SNP linkage disequilibrium (LD) data will tend to underestimate *N*_e_ because of physical linkage between many of the SNPs in the data set (Waples et al., 2016). Nevertheless, taking this into account, there is still cause for concern that the most recent *N*_e_ values modelled for the KY92 and KY12 population samples are below the critical *N*_e_ threshold of 100 recommended by Meuwissen (2009) for long-term viability of discrete livestock breeds and populations.

### 3.7. Genomic Relationship and Analysis of Inbreeding

Supplementary Table 1 shows a genomic relationship matrix in terms of genotype IBS for the genome-wide SNP data generated for individual animals in the KY92 and KY12 population samples. Close genomic relationship between individual animals sampled from the same herd is evident in the SNP genotype IBS values between samples. In addition, the relatively low genomic relationship between the KY12_06 and KY12_58 outlier samples (**Figures 4–7**) and the rest of the Kerry cattle sampled is also evident in Supplementary Table 1. These data emphasise the value of intrapopulation genomic relationship values for identifying animals that, from a conservation genetics standpoint, should be of lower priority. They also highlight the potential of genome-wide SNP data for providing a systematic approach to prioritising males and females with minimum genomic relationship for breeding to minimize loss of genetic diversity and maintain or increase *N*_e_ (Gandini et al., 2004; Meuwissen, 2009; de Cara et al., 2011; de Cara et al., 2013).

Genome-wide autozygosity estimated from SNP data using runs of homozygosity (ROH) and the *F*_ROH_ statistic are visualised in **Figure 9** for individual animals from the KY92 and KY12 population samples and a range of European comparator breeds. Additional summary ROH data is provided in Supplementary Table 3 and also Supplementary Figure 2, which reveals marked inter-population differences in ROH length and demonstrates that the SNP density of the Illumina^®^ Bovine SNP50 BeadChip is too low to reliably capture ROH below 5 Mb in length, an observation previously reported by Purfield and colleagues (2012).

**FIGURE 9.**
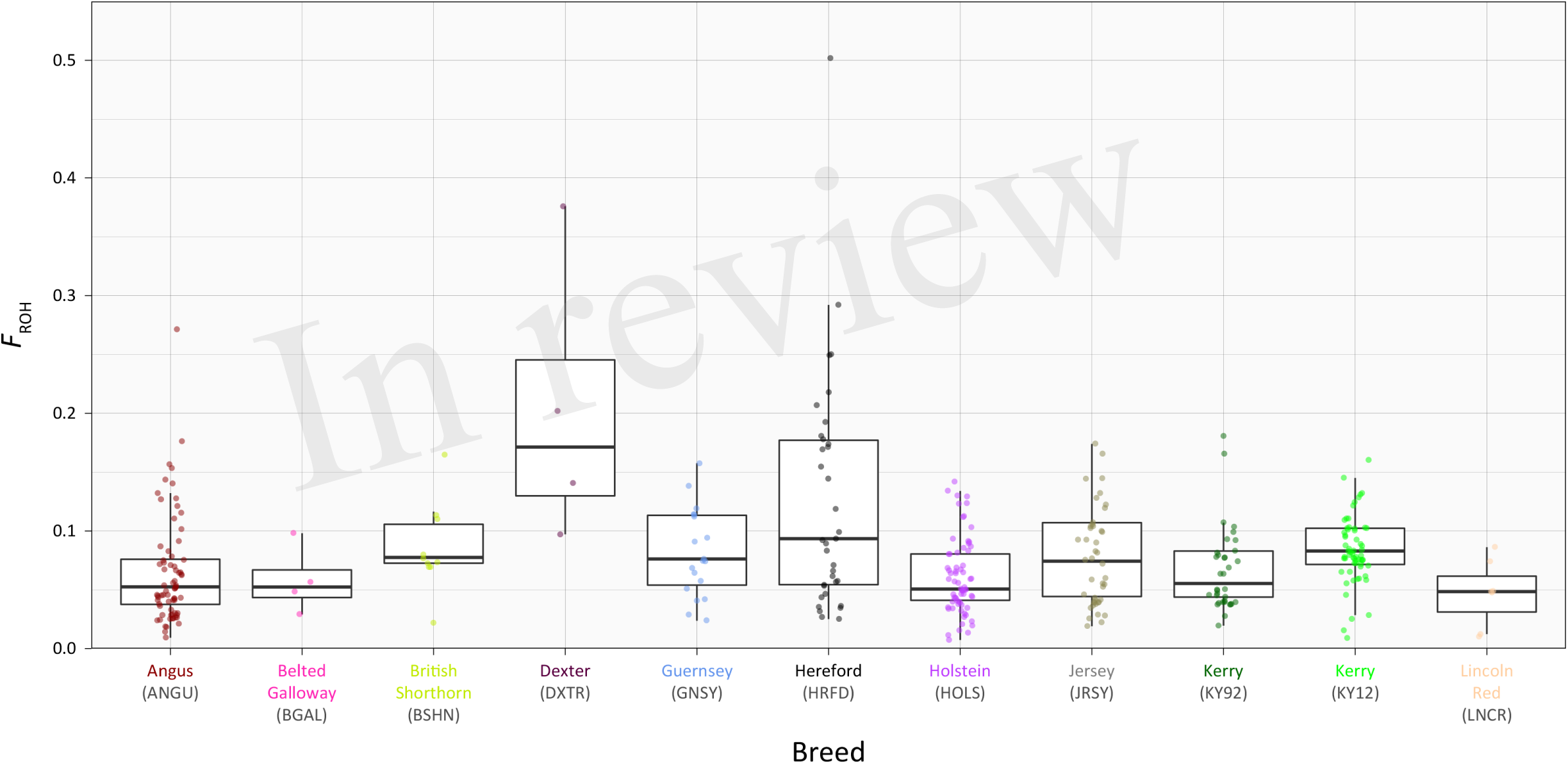
Tukey box plots showing distribution of *F*_ROH_ values estimated with genome-wide SNP data for the KY92 and KY12 populations and nine comparator heritage and production cattle breeds.

There is significant variation in *F*_ROH_ values among individual animals and between breeds and populations. The non-parametric Wilcoxon rank sum test was performed on *F*_ROH_ distributions for all pairwise population/breed comparisons with application of the Bonferroni correction *P* value adjustment for multiple statistical tests (Supplementary Table 4). This analysis demonstrated that the KY12 population sample exhibited a significantly higher mean *F*_ROH_ value than the KY92 population sample (*P*_adjust_ = 0.0340). This is important from a conservation genetics perspective, indicating that genome-wide autozygosity, which is highly correlated with conventional pedigree-based estimates of inbreeding (*F*_PED_) for cattle (Purfield et al., 2012; Ferenčaković et al., 2013; Martikainen et al., 2017), has increased for the Kerry cattle population in the 20 years between sampling of the KY92 and KY12 populations.

The importance of understanding and quantifying genome-wide autozygosity for genetic conservation purposes has recently been highlighted through correlation of *F*_ROH_ with inbreeding depression for a range of production traits in domestic cattle (Bjelland et al., 2013; Pryce et al., 2014; Kim et al., 2015). Importantly, *F*_ROH_ has also been shown to correlate with inbreeding depression for bovine fertility traits in both males (Ferencakovic et al., 2017) and females (Kim et al., 2015; Martikainen et al., 2017). Finally, according to basic population genetic principles, recent inbreeding captured by *F*_ROH_ will lead to recessive deleterious genomic variants emerging at a population level— a phenomenon that has been studied in both humans and cattle (Szpiech et al., 2013; Zhang et al., 2015).

### 3.8. Genome-wide Signatures of Selection in the Kerry Cattle Breed

The results of the genome-wide scan for signatures of selection using the CSS method in the Kerry cattle breed are shown in **Figure 10**. Six distinct selection signatures were detected on BTA9, BTA12, BTA16, BTA17, BTA19 and BTA28. A total of 178 genes were located within the genomic ranges ± 1.0 Mb of selection peaks and 32 of these genes were located within the boundaries of a selection peak. Supplementary Table 5 provides detailed information for these 178 genes.

**FIGURE 10.**
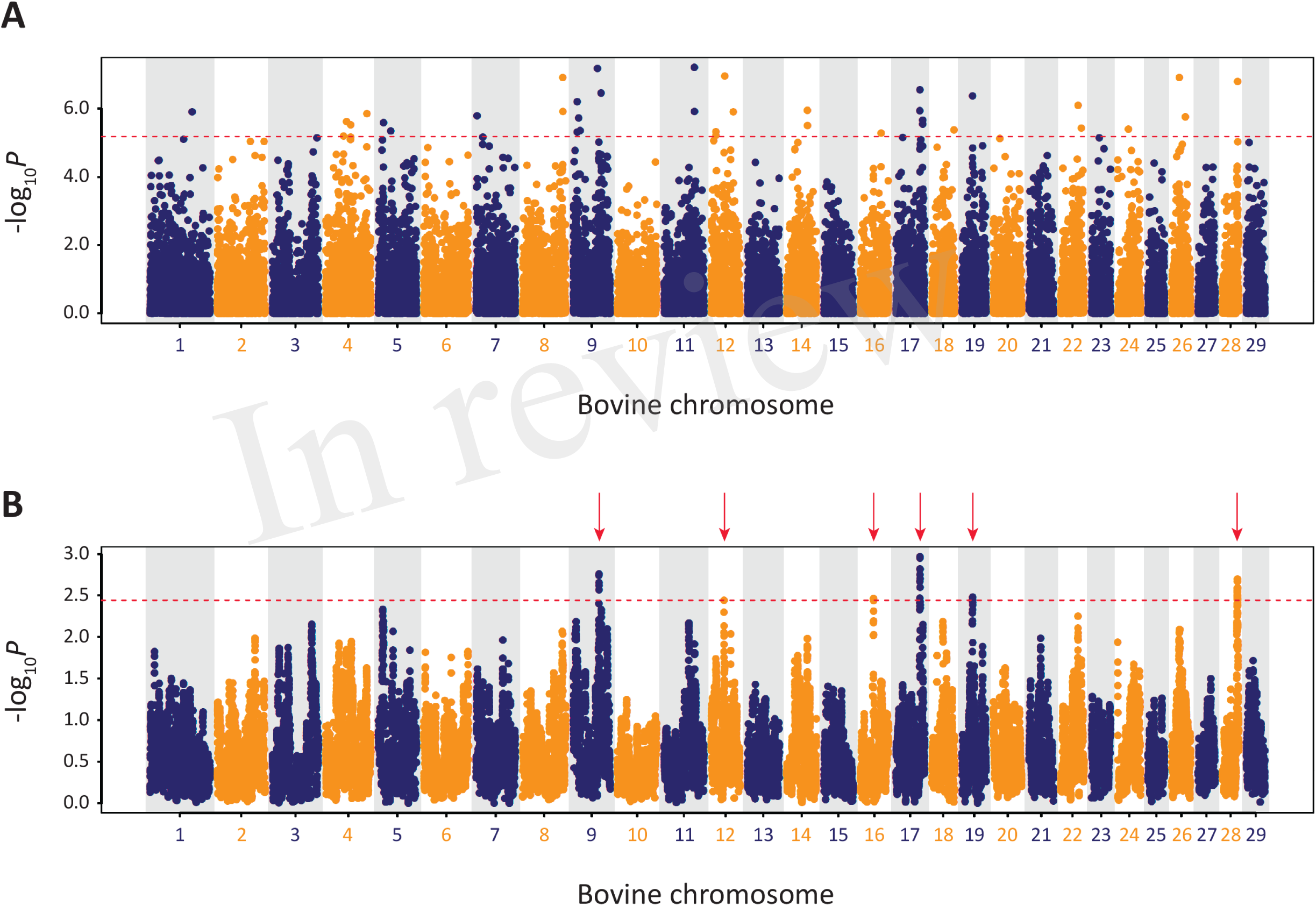
Manhattan plots of composite selection signal (CSS) results for Kerry cattle (*n* = 98) contrasted with EU cattle (*n* = 102). A) Unsmoothed results. B) Smoothed results obtained by averaging CSS of SNPs within each 1 Mb window. Red dotted line on each plot denotes the genome-wide 0.1% threshold for the empirical CSS scores. Red vertical arrows indicate selection peaks detected on BTA09, BTA12, BTA16, BTA17, BTA19 and BTA28.

A single gene was located within the BTA9 selection peak—the phosphodiesterase 7B gene (*PDE7B*), which has been associated with neurobiological processes (de Gortari and Mengod, 2010) and has been previously linked to genetic changes associated with dog (*Canis lupus familiaris*) domestication and behaviour (Freedman et al., 2016). A single gene was also located within the BTA16 selection peak—the dorsal inhibitory axon guidance protein gene (*DRAXIN*), which encodes a protein that regulates axon guidance, neural circuit formation and vertebrate brain development (Islam et al., 2009; Shinmyo et al., 2015). Twenty-four genes were located within the BTA17 selection peak, including *BICDL1*, *RAB35* and *RNF10*, which have been associated with neurobiology and brain development (Hoshikawa et al., 2008; Schlager et al., 2010; Villarroel-Campos et al., 2016) and *SIRT4* and *COQ5* that function in cellular metabolism (Kawamukai, 2015; Elkhwanky and Hakkola, 2017). Six genes were located within the BTA28 selection peak, including, most notably, the Rho GTPase activating protein 22 gene (*ARHGAP22*), which has recently been associated with bovine fertility as an mRNA expression biomarker for oocyte competence in cumulus cells (Melo et al., 2017).

To obtain a broader perspective on natural and artificial selection acting at a population level on the Kerry cattle genome, a functional gene set enrichment approach (GSEA) was taken using IPA with the 178 genes located within ± 1.0 Mb of each selection peak (Supplementary Table 5). Of these 178 genes, 141 could be mapped to the IPA knowledgebase and the summary results for the IPA *Physiological System Development and Function* category are shown in Supplementary Table 6, revealing an enrichment of biological processes associated with nervous system development and behaviour.

### 3.9. Genomics, Genetic Distinctiveness and Microevolution of Kerry Cattle: Implications for Breed Management and Genetic Conservation

The genome-wide phylogenetic and population genetic analyses detailed here demonstrate that Kerry cattle represent an important farm animal genetic resource, befitting the breed’s status as a livestock population with a unique history of adaptation to the climate and physical geography of southwest Ireland at the edge of Western Europe. Notably, from a genetic conservation and breed management perspective, high-resolution comparative PCA (**Figures 4** and **5**) and genetic clustering results (**Figures 6** and **7**) demonstrate that Kerry cattle are markedly distinct from other British and European cattle populations. This observation may also be placed in the context of recent paleogenomic studies that have detected ancient gene flow from wild British aurochs (*B. primigenius*) into the ancestors of present-day Kerry cattle (Orlando, 2015; Park et al., 2015; Upadhyay et al., 2017).

The current genetic status of the Kerry cattle population is underlined by analyses of genetic effective population size (*N*_e_) and inbreeding using genome-wide SNP data. As shown in **Table 2**, genome-wide observed heterozygosity (*H*_o_) is relatively high in the KY92 and K12 population samples, particularly for endangered heritage cattle breeds. However, it has been long recognised that monitoring *N*_e_ is a more important tool for rational breed management and long-term conservation of endangered livestock populations (Notter, 1999; Gandini et al., 2004; Biscarini et al., 2015). As shown in **Figure 8** and Supplementary Table 2, the Kerry cattle population has a recent demographic trend of *N*_e_ decline, to the point where the most recent modelled *N*_e_ values are below the recommended threshold for sustainable breed management and conservation (Meuwissen, 2009). There is also cause for concern that genomic inbreeding estimated using genome-wide autozygosity (*F*_ROH_) and visualised in **Figure 9** has increased significantly in the 20-year period between the sampling of the KY92 and KY12 Kerry cattle populations.

In a more positive light, as shown in the present study, detection of discrete signatures of selection using the relatively low-density Illumina^®^ Bovine SNP50 BeadChip is encouraging for wider studies of genome-wide microevolution in endangered heritage livestock populations. Based on our results, for example, future surveys of Kerry cattle that use higher-density SNP array platforms and ultimately whole-genome sequence data will provide exquisitely detailed information on the genomic regions and associated polygenic production, health, fertility and behavioural traits shaped, over many centuries, by the agro-ecology and pre-industrial farming systems of southwest Ireland.

## 4. Conflict of Interest

The authors declare that the research was conducted in the absence of any commercial or financial relationships that could be construed as a potential conflict of interest.

## 5. Ethics Statement

With the exception of Kerry cattle sampled during 2011-12, all samples and data was obtained from previously published scientific studies. The re-use of these samples and data is consistent with the 3Rs principles on replacement, refinement and reduction of animals in research (www.nc3rs.org.uk/the-3rs). For the 2011-12 Kerry cattle, population owners’ consent to sample DNA for research was obtained and individual owners conducted sampling of animals using non-invasive nasal swabs. In this regard, scientific animal protection in Ireland is subject to European Union Directive 2010/63/EU, which states that the Directive does not apply to “practices not likely to cause pain, suffering, distress or lasting harm equivalent to, or higher than, that caused by the introduction of a needle in accordance with good veterinary practice”.

## 6. Author Contributions

DEM, DAM, AGF and JFK conceived and designed the project; DEM, IWR, DAM, AGF and JFK organised sample collection and genotyping; SB, GM, IWR, DAM, SDEP, CNC, IASR and DEM performed the analyses; SB and DEM wrote the manuscript and all authors reviewed and approved the final manuscript.

## 7. Funding

This work was supported by Department of Agriculture, Food and the Marine (DAFM) funding under the Genetic Resources for Food and Agriculture scheme (grant no: 10/GR/06); an Investigator Programme Grant from Science Foundation Ireland (SFI/08/IN.1/B2038); a Research Stimulus Grant from DAFM (RSF 06 406); a European Union Framework 7 Project Grant (KBBE-211602-MACROSYS); the Brazilian Science Without Borders Programme (CAPES grant no. BEX-13070-13-4) and the UCD MSc Programme in Evolutionary Biology.

## 8. Acknowledgements

The authors wish to express their gratitude to the Kerry Cattle Society (www.kerrycattle.ie) for facilitating and encouraging this project. In particular, we thank Rosemary and Jeremy Hill, Matthew English Hayden and the Irish Cattle Breeding Federation (ICBF – www.icbf.com) for expert guidance and assistance with animal sourcing and DNA sampling. We are also grateful to Professor Dan Bradley (Smurfit Institute of Genetics, Trinity College Dublin) for access to DNA sample archives. In addition, we thank all of the Kerry cattle owners who provided access to animals, samples and pedigree information. Finally, we thank Weatherbys Scientific for provision of SNP array genotyping services (www.weatherbysscientific.com).

